# Prediction of gene expression from regulatory sequence composition enhances transcriptome-wide association studies

**DOI:** 10.1101/2021.05.11.443571

**Authors:** Federico Marotta, Reza Mozafari, Elena Grassi, Alessandro Lussana, Elisa Mariella, Paolo Provero

## Abstract

Transcriptome-wide association studies (TWAS) can prioritize trait-associated genes by finding correlations between a trait and the genetically regulated component of gene expression. A basic ingredient of a TWAS is a regression model, typically trained in an external reference data set, used to impute the genetically-regulated expression. We devised a model that improves the accuracy of the imputation by using, as predictors, not the genotypes directly but rather the sequence composition of the proximal gene regulatory region, expressed as its profile of affinities for a set of position weight matrices. When trained on 48 tissues from GTEx, the regression model showed improved performance compared with models regressing expression directly on the genotype. We imputed the expression levels in genotyped individuals from the ADNI data set, and used the imputed expression to perform a TWAS. We also developed a method to perform the TWAS based on summary statistics from genome-wide association studies, and applied it to 11 complex traits from the UK Biobank. The greater accuracy in the prediction of gene expression allowed us to report hundreds of new gene-phenotype association candidates.

## 1 Introduction

Genome-wide association studies (GWAS) have revealed thousands of associations between genetic variants and complex traits or diseases [1, 2]. Most of such GWAS hits lie outside coding regions, and are likely to affect the phenotype by altering gene regulation, a view supported by various strands of evidence [3, 4, 5]. However the regulatory effects of a non-coding variant, and even its target genes, are difficult to predict. This fact, combined with linkage disequilibrium which makes it difficult to identify the truly causal variants among those showing statistically significant associations, makes the interpretation of GWAS hits and their eventual exploitation in devising new approaches to treatment extremely challenging.

Recently, new approaches have been developed to tackle these issues and, specifically, to derive gene/trait associations, rather than variant/trait associations, from GWAS data, based on intermediate molecular phenotypes. In these methods genetic variants are first associated, using a reference data set, to a suitable molecular phenotype such as mRNA expression [6, 7], splicing isoform abundance [8], or protein expression [9]. A predictive model is generated from the reference data and used to impute the intermediate molecular phenotype in the individuals assayed by the GWAS, then associations are sought between the imputed molecular phenotype and the trait of interest.

Based on the putative regulatory effect of GWAS hits, gene expression is a natural choice of intermediate phenotype, and was indeed used in the first such studies, which were thus called transcriptome-wide association studies (TWAS) [6]. To train the predictive model of gene expression, TWAS can rely on large dataset, such as GTEx [10], where genotypes and gene expression data in various tissues are available for hundreds of individuals. TWAS can also be perfomed when only GWAS summary statistics are available, by combining in a statistically appropriate way the expression weights with the GWAS Z-scores [7, 11].

As the final goal of a TWAS is the discovery of associations between a phenotype and the genetically-regulated component of gene expression, a reliable method to infer the genetically-regulated component of expression is needed. The common way to tackle this problem is to build, for each gene, a regression model where the outcome is the expression and the predictors are the 0/1/2-encoded genotypes for all the markers falling in a predefined window around the gene [6, 12]. Since the predictive model uses only DNA-derived features as predictors, the differences in expression that it captures represent the genetically-determined variance in gene expression.

Since the seminal paper by Gamazon *et al.* [6], most predictive models applied to TWAS have used, as predictors, a direct 0/1/2 encoding of the genotypes, and the differences between the tools proposed so far amount to the level of sophistication of the statistical learning model employed. For instance, PrediXcan [6] uses elastic net; FUSION [7] relies on an ensemble of models, including a best linear unbiased predictor (BLUP) and a Bayesian sparse linear mixed model (BSLMM); and TIGAR [13] introduces a complex hierarchical Bayesian model. Of notice, HAPLEXER [14] is different in that it uses signature tract sets (*i.e.* sets of haplotypes shared between individuals) instead of genotypes, and TF2Exp [15] uses the predicted differences in the binding of transcription factors to their target sites.

In this work, we present a new strategy, whereby gene expression is predicted from the sequence composition of a gene’s proximal regulatory region. Sequence composition is usually expressed as the frequency spectrum of k-mers; here, based on our previous work on eQTLs [16], we chose to parametrize sequence composition as the spectrum of total bindig affinity (TBA) for a set of position weight matrices (PWMs) associated to transcription factors (TFs), as these are the DNA motifs most likely to impact gene expression. A gene’s regulatory sequence is thus represented by a TBA profile, which is used as regressor in predicting the variation of gene expression across individuals.

We show that this strategy outperforms the previously proposed ones, even using a relatively simple regression model such as Ridge regression, and even though we consider only proximal regulatory regions as opposed to large windows around each gene. We also describe how our strategy can be used when only GWAS summary statistics are available, albeit in an approximate form. Our models were trained in 46 tissues and 2 cell lines from GTEx (in the following collectively denoted as “tissues”) and applied to perform genotype-level TWAS for 43 Alzheimer’s-related phenotypes measured in the Alzheimer Disease Neuroimaging Initiative (ADNI), and summary-level studies for 11 complex traits in the UK Biobank, allowing us to report hundreds of new genes potentially associated to complex traits or diseases. We also developed and made available a convenient suite of scripts to run these analysis (Section 4.11).

## 2 Results

### 2.1 Prediction of gene expression from TBA profiles

In our approach variation across individuals of the expression of a gene in a given tissue is modeled as depending on the TBA for a set of PWMs of the proximal regulatory region of the gene. The TBA, inspired by a statistical-mechanical model [17], integrates the signal given by the PWM along all the possible binding sites in the regulatory region. We have previously shown that it can be used to effectively encode differences in gene regulation among genes [18], species [19], and individuals [16]. Therefore, our model assumes that genetically determined differences among individuals in the expression of a gene in a given tissue are mediated by the differences in the TBA profiles of the gene’s regulatory region.

For each gene, we defined its regulatory region as the union of all the sequences extending from 1500 base pairs (bps) upstream to 500 bps downstream of the transcription start site (TSS) of all the annotated transcripts; then, given a library of PWMs and a reference panel of individuals with their genomes sequenced, we can compute the TBA for each PWM of the regulatory region of each gene in each individual. The TBA values of the reference individuals can then be used, instead of the genotypes, as independent variables in a penalized regression model where the dependent variable is gene expression. Preliminary analysis suggested that Ridge regression outperformed both least absolute shrinkage and selection operator (Lasso) and elastic net in predicting expression, thus Ridge regression was used to build all the predictive models. The genetically regulated component of expression has been previously abbreviated with GReX [6]; since our model finds the genetic component mediated by the total binding affinity, we called our tool TReX, from “TBA-Regulated eXpression.” We also refer to the imputed expression as 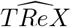.

Using the subset of white individuals from the GTEx data set [10] as a reference panel and Hocomoco [20] for the position weight matrices, we computed the affinity for 640 PWMs and trained our model to predict the expression of both coding and non-coding genes in 48 tissues. The number of genes in each tissue was close to 40,000, whereas the number of individuals ranged from 70 for substantia nigra to 421 for skeletal muscle. We thus trained almost two million models to predict the expression of a gene in a given tissue starting from the DNA sequence of its regulatory region in a given individual.

As it is customary [6, 13, 15], in order to assess the predictive performance of our Ridge model andcompare it with that of other tools, we used 5-fold cross-validation, assessing the performance by the *R*^2^ value, defined as the squared correlation coefficient between true and predicted expression values in the validation fold, averaged over the 5 folds. We compared the cross-validation *R*^2^ achieved by TReX with those achieved on the same data by TIGAR [13], which, to the best of our knowledge, is currently the best performing prediction tool based on the same performance metric and the same data.

Table 1 shows some statistics related to the distribution of 5-fold cross-validated *R*^2^s obtained with the two tools in EBV-transformed lymphocytes (Fig. 1 (**a** and **b**)) and brain cortex (Suppl. Fig. 1). The difference in performance was statistically significant (*P* < 2.2*e* − 16, paired Wilcoxon test on *R*^2^, in both tissues). Cross-validated *R*^2^ values for all GTEx tissues are reported in Suppl. Tab. 1. The improvement in performance brought by TReX was significant in both protein coding and non-coding genes considered separately, but the increase in cross-validated *R*^2^ was significantly higher for protein-coding genes (*P* < 2.2*e* − 16 in both tissues; Mann-Whitney U test).

**Table 1:**
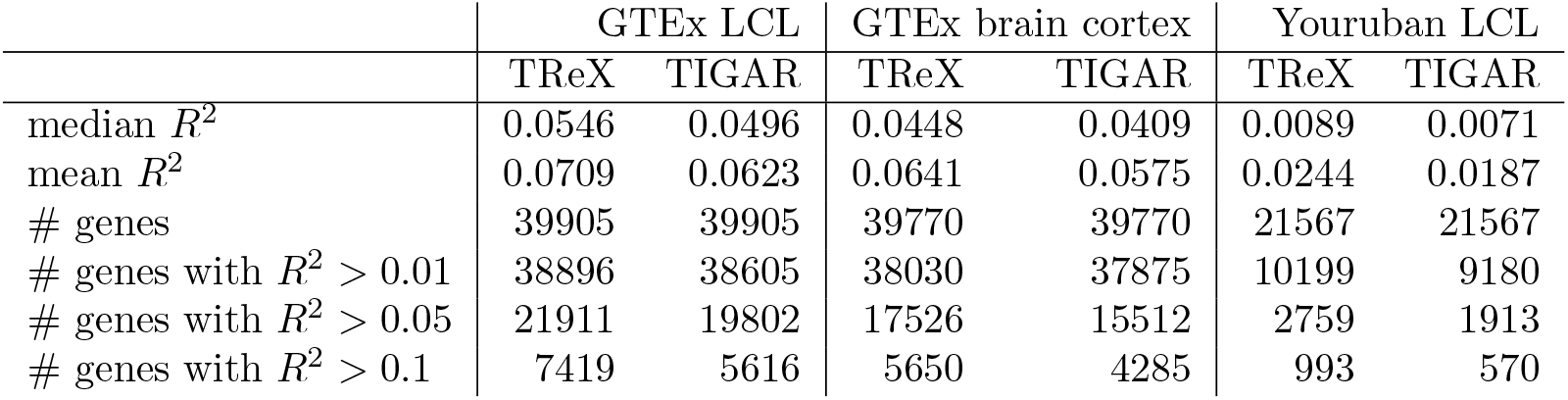
*R*^2^ distribution for two GTEx tissues and an external dataset representing a different ethnicity. *R*^2^ is defined as the square of the correlation between predicted and actual values (averaged over 5 cross-validation folds in the GTEx tissues).

|  | GTEx LCL |  | GTEx brain cortex |  | Yourubian LCL |  |
| --- | --- | --- | --- | --- | --- | --- |
|  | TReX | TIGAR | TReX | TIGAR | TReX | TIGAR |
| median $R^2$ | 0.0546 | 0.0496 | 0.0448 | 0.0409 | 0.0089 | 0.0071 |
| mean $R^2$ | 0.0709 | 0.0623 | 0.0641 | 0.0575 | 0.0244 | 0.0187 |
| # genes | 39905 | 39905 | 39770 | 39770 | 21567 | 21567 |
| # genes with $R^2 > 0.01$ | 38896 | 38605 | 38030 | 37875 | 10199 | 9180 |
| # genes with $R^2 > 0.05$ | 21911 | 19802 | 17526 | 15512 | 2759 | 1913 |
| # genes with $R^2 > 0.1$ | 7419 | 5616 | 5650 | 4285 | 993 | 570 |

**Figure 1:**
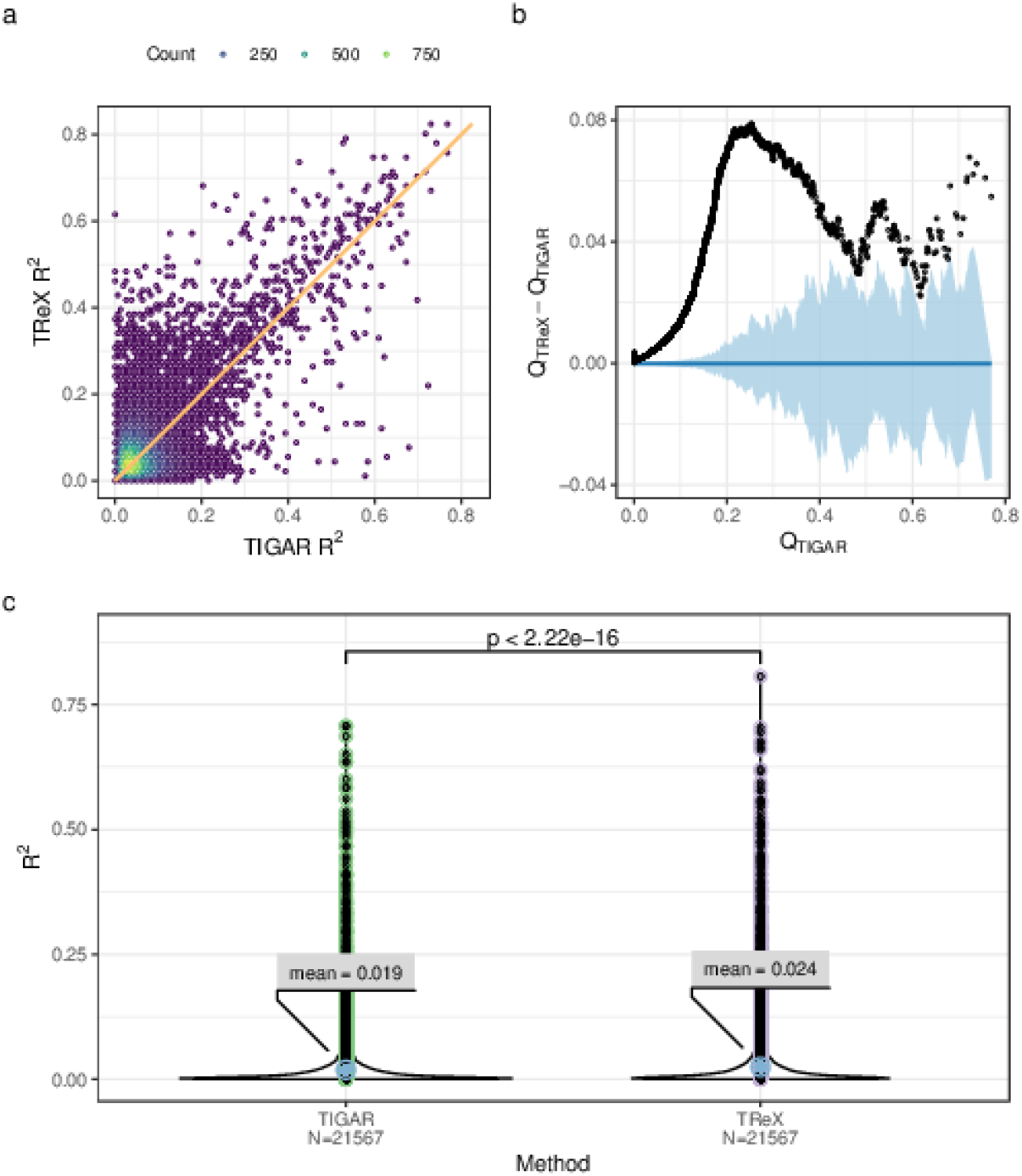
Predictive performance of various models. **a.** Scatter plot of the 5-fold cross-validation *R*^2^ for TIGAR and TReX on EBV-transformed lymphocytes; the plot shows 39905 genes. In principle each point represents a gene, but in order to reduce overplotting we associated potentially many genes that fell approximately in the same region to a single point; the color of a point denotes the number of genes associated to it. The orange line is the identity line. **b.** Detrended QQ-plot of the 5-fold cross-validation *R*^2^ for TIGAR and TReX on EBV-transformed lymphocytes. The x axis represents the quantiles of TIGAR’s *R*^2^, while on the y axis there are the differences between corresponding quantiles from TReX’s and TIGAR’s *R*^2^. The confidence interval (light blue) was generated from 100 bootstrap replicates from the distribution of TIGAR’s *R*^2^. **c.** Predictive performance in an external data set. Distribution of the squared correlation between the true and predicted expression values in 89 Yoruban individuals from the GEUVADIS data set (21567 genes common to both methods).

The improvement in predictive performance was not limited to cross-validated samples but extended to the prediction of gene expression in samples from a different dataset and population: When we used our model to predict gene expression in LCLs in a sample of 89 Yoruba individuals from the Geuvadis dataset, we obtained again *R*^2^ values significantly higher that those obtained with TIGAR (Fig. 1 (**c**) and Table 1). About half the genes with *R*^2^ > 0.1 according to TIGAR also have *R*^2^ > 0.1 according to TReX (Suppl. Fig. 2), suggesting that the genes on which the two methods are most effective do not coincide, so that integrating the two could provide even greater predictive power.

A direct comparison with TF2Exp [15], which like TReX predicts expression variation based on predicted differences in transcription factor affinity, is made problematic by the fact that TF2Exp was trained on the Geuvadis LCL dataset rather than of GTEx tissues. Qualitatively, both the percentage of genes with cross-validated *R*^2^ > 0.05 (21911/40056 = 54.7%) and the mean *R*^2^ (0.0709) of TReX are higher than the corresponding values reported [15] for TF2Exp (respectively 20.1% and 0.049). More generally, TF2exp uses cell-type specific epigenetic information which is not readily available for most of the GTEx tissues (see also Discussion).

Therefore encoding the genotype as a profile of total binding affinities for PWMs led to a significant improvement of the prediction, even if only genetic variants within a region of length ~ 2, 000 were used to compute such profiles. It would be tempting to interpret these results as reflecting the underlying biology of gene regulation, by stating that the TBA profiles capture the genetic variation that is responsible for inter-individual differences in TF binding, which is in turn responsible for differences in gene expression. However, models trained with scrambled versions of the position weight matrices, which are unlikely to reflect the binding preferences of any TF, gave results very similar to those obtained with the original PWMs (Suppl. Fig. 3). Therefore, the reason for the improvement in performance must be that affinity profiles for PWMs provide an efficient encoding of sequence composition allowing non-linear effects of the genotype on expression to be taken into account (see also Discussion).

### 2.2 Genotype-level TWAS

We computed the total binding affinities of the Hocomoco PWMs for the proximal regulatory regions of all the genes in 735 white individuals from the ADNI data set for which the genotypes and 43 Alzheimer’s disease (AD)-related phenotypes were available. Then, using the models previously trained on the 48 GTEx tissues, we imputed the expression in these individuals from their binding affinities, and found associations between the predicted expression and each of the phenotypes using linear or logistic regression, according to whether the phenotype was quantitative or categorical. In all regressions we included the same set of covariates (see Section 4.6). Two of the phenotypes were categorical and pertained to the diagnosis at the baseline visit, while the 41 quantitative phenotypes included volumetric measures of brain regions, quantifications of biomarkers such as TAU protein and amyloid *β*, and results of cognitive tests such as the Preclinical Alzheimer Cognitive Composite (PACC), the Alzheimer’s Disease Assessment Scale-Cognitive (ADAS), or the Rey’s Auditory Verbal Learning Test (RAVLT). The associations were investigated on a gene-by-gene basis, therefore, in total, we tested about 10 million associations. Given the high correlation among the phenotypes considered and among the predicted gene expression in different tissues, we implemented an independent Bonferroni correction for each tissue/phenotype pair, finding 179 significant associations involving 84 genes, 47 tissues, and 43 phenotypes (Suppl. Table 2 and Section 4.12). The strongest associations (Figure 2 **c, d**) appeared to be biologically meaningful and supported by the scientific literature. As it will be shown in the next section, we exploited this TWAS on ADNI also to compare the results of the genotype- and summary-level TWAS on the same data.

**Figure 2:**
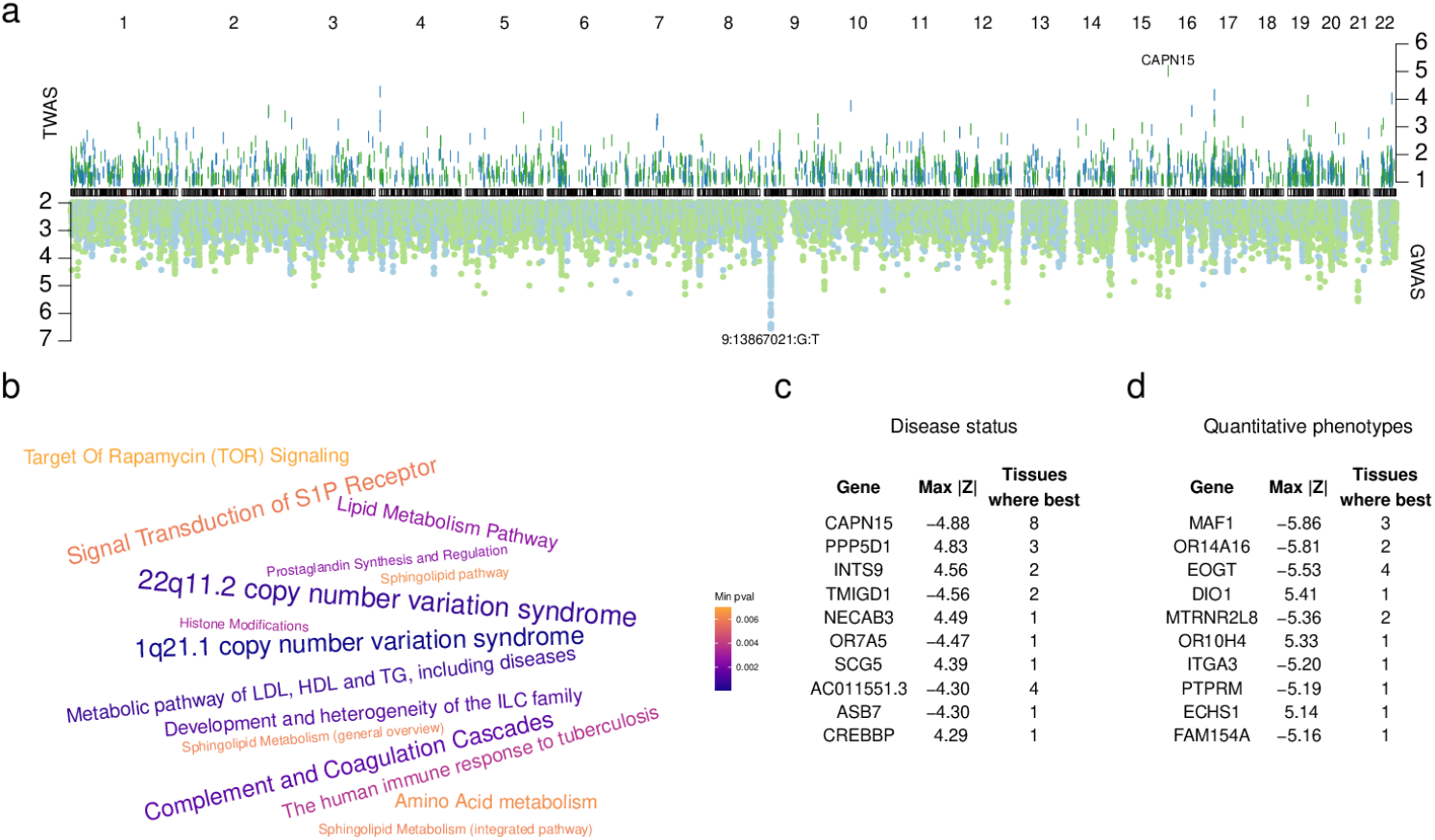
Transcriptome-wide association study on the ADNI data set. **a.** Double Manhattan plot for disease status. The Z-scores of allgenetic markers (bottom panel) and all genes with cross-validated *R*^2^ > 0.1 (top panel) are shown as a function of their genomic location. For the TWAS, the Ridge model for the imputation of gene expression was trained on the hypothalamus GTEx data. Blue (green) points or bars indicate positive (negative) association with disease status encoded as 0 for controls and 1 for patients diagnosed as Alzheimer’s disease, early/late mild cognitive impairment, or significant memory concern (see 4.5). **b.** Top enriched pathways for the TWAS on the “Ecog” quantitative phenotype. The size of each term is proportional to the number of tissues in which the term is enriched with an adjusted p-value smaller than 0.05, ranging from 1, (*e.g.* Histone Modifications), to 7 (22q11.2 copy number variation syndrome). The color of each term represents the smallest adjusted enrichment p-value. **c, d.** Top genes associated to disease status or a quantitative phenotype. Associations between the disease status and the predicted expressions were found through logistic regression; disease status was encoded in two different ways as described in 4.5. Associations with quantitative phenotypes were assessed through linear regression. The tables report, for each gene, the maximum Z-score across all 48 tissues, and the number of tissues where that gene had the best Z-score for at least one of the phenotypes.

The strongest association with disease status among protein-coding genes was shown by the expression of CAPN15 in the colon transverse. CAPN15 was also the top scoring gene for the association with disease status in seven other tissues (hypothalamus, esophagus mucosa, ovary, pancreas, prostate, skin - not sun exposed suprapubic, skin - sun exposed lower leg; Figure 2 **c**). The calpains, to which this gene belongs, are a family of cysteine proteases whose hyperactivation has been previously related to Alzheimer’s disease [21]; furthermore, a recent preprint describes how the knockout of CAPN15 in mice impairs brain development and plasticity [22]. Interestingly, CAPN15 has not been previously associated to AD through a GWAS according to the EBI GWAS catalog, showing that our method can find new gene/disease associations.

PPP5D1, a protein-serine/threonine phosphatases, was also strongly associated to disease status in multiple tissues. This gene is poorly characterized in the literature, but in the GWAS catalog it is associated to late-onset AD. INTS9 likely plays a role in small nuclear RNA processing [23], and was found differentially expressed in AD patients [24]. Remarkably, we also found NECAB3 among the top-scoring genes: This calcium binding protein inhibits the association of X11L with amyloid precursor protein and abolishes the suppression of beta-amyloid production by X11L [25, 26]. We could not find any reliable reference describing an association between TMIGD1 and Alzheimer’s disease, but it was associated with Crohn’s disease [27], and in turn inflammatory bowel disease seems to be related to Alzheimer’s [28].

To further demonstrate our results on the ADNI data set, we chose a particular tissue, the hypothalamus, and plotted both the GWAS and the TWAS for disease status together (Figure 2 **a**). Notably, many TWAS hits do not correspond to a GWAS peak.

The results for the quantitative phenotypes were analyzed together by finding, for each tissue, the gene with the strongest association considering all 41 phenotypes together, thus obtaining a list of one gene per tissue. The top 10 genes by maximum absolute Z-score together with the number of tissues in which they show the strongest association are shown in Figure 2 **d**.

The genetic component of MAF1 expression in three tissues (brain cortex, cervical spine, and gastroesophageal junction) is strongly and negatively associated to the volume of the hippocampus; consistently, it was recently reported that MAF1 inhibits dendritic morphogenesis and the growth of dendritic spines, and affects learning and memory in mice [29]. Notably, the association between MAF1 expression in the hippocampus itself and hippocampal volume was not significant. This apparent contradiction could be explained either by an effect of the phenotype on gene expression (which, for continuous phenotypes, would be present also in the reference dataset; see Supplementary Note, Sec.1.2) or by a genetic variant pleiotropically affecting both expression and phenotype.

Also the imputed expression of EOGT significantly correlates with several Alzheimer-related phenotypes, although this gene has not been implicated in AD by GWAS studies. The role of EOGT is to attach sugar moieties to extracellularly-secreted or membrane proteins [30]. Glycosylation is known to regulate the cleavage of the amyloid precursor protein (APP) [31], and has also been implicated in many forms of neurodegeneration [32, 33].

To the best of our knowledge, all attempts to find associations between polymor-phisims in the DIO1 gene (whose product, D1, can activate or inactivate the thyroid hormone) and cognitive functions so far have failed [34, 35], despite the fact that thyroid function is linked to Alzheimer’s disease [36]. Here, we discovered a positive, relatively strong association between the 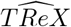 of DIO1 in the adrenal gland and the results of the EcogSPDivatt test.

So far we have only reported associations with protein-coding genes, but we actually found many associations with pseudogenes or other non-coding genes. These associations are more difficult to explain because the genes involved are not well-documented in the literature.

We also investigated whether relevant biological pathways are enriched among the top-scoring genes. For each tissue-phenotype pair, we selected the genes with Z-score greater than 3 in absolute value, and analyzed them for enriched pathways from the WikiPathways database. After pooling together all the four phenotypes pertaining to RAVLT scores, we considered the pathways with an enrichment adjusted p-value smaller than 0.05, and counted in how many different tissue each pathway appeared; the results are shown in Figure 2 **b**.

In the literature, we could find evidence for the implication of all these terms to cognitive disorders. For instance, 22q11.2 CNV syndrome is known to alter the structure of the brain [37] and lead to cognitive decline [38]. Another CNV syndrome, 1q21.1, was detected in Alzheimer’s disease patients [39]. Inflammation has been long known to lead to several diseases, including neurodegenerative ones; thus, it is not surprising here to find pathways related to innate lymphoid cells (ILC), complement and coagulation [40, 41]. We also found many terms related to lipids, which is, again, not surprising, as lipids play many fundamental roles in the brain: from the processing of amyloid precursor protein (APP) to receptor signaling, from myelination to membrane remodeling, from oxidation to energy balance; the implication of lipids in Alzheimer’s disease has been reported many times [42, 43, 44].

In conclusion, although multiple testing correction was performed separately for each tissue/phenotype pair, inspection of the significant associations seems to confirm the validity of the approach. To exploit GWAS data derived from much greater samples, but available only as summary results, we developed a strategy to perform a summary-level TWAS using our affinity-based weights, and applied it to the UK Biobank, where summary GWAS results based on hundreds of thousand of individuals are available.

### 2.3 Summary-level TWAS

As we mentioned in the previous section, the sample size in data sets where the genotype of each individual is available is still very limited, either for privacy concerns or logistic reasons [7]. Much more commonly, only the summary statistics of a GWAS are available; Gusev *et al.* [7] developed a methos to perform a TWAS using only a model trained in a reference data set and the GWAS summary statistics. However, their method was developed to work with the 0/1/2-encoded genotypes rather than with genotypes summarized as TBA profiles or otherwise. Here, we extended their method so that it could be applied also when the expression-predicting model is trained on TBA profiles (see Section 4.9). Briefly, we computed the change in TBA determined by each SNP, and combined it with the weights learned during the training phase to find the change in expression determined by each SNP through the induced changes TBA. Thus, we reduced to the original case and could apply the association test developed by Gusev *et al.* in an approximate version. We performed a summary-level TWAS on the same ADNI data used for the genotype-level TWAS described above, and we found the results of summary-leval and genotype-level TBA-based TWAS to be highly correlated (Figure 3 **a**).

**Figure 3:**
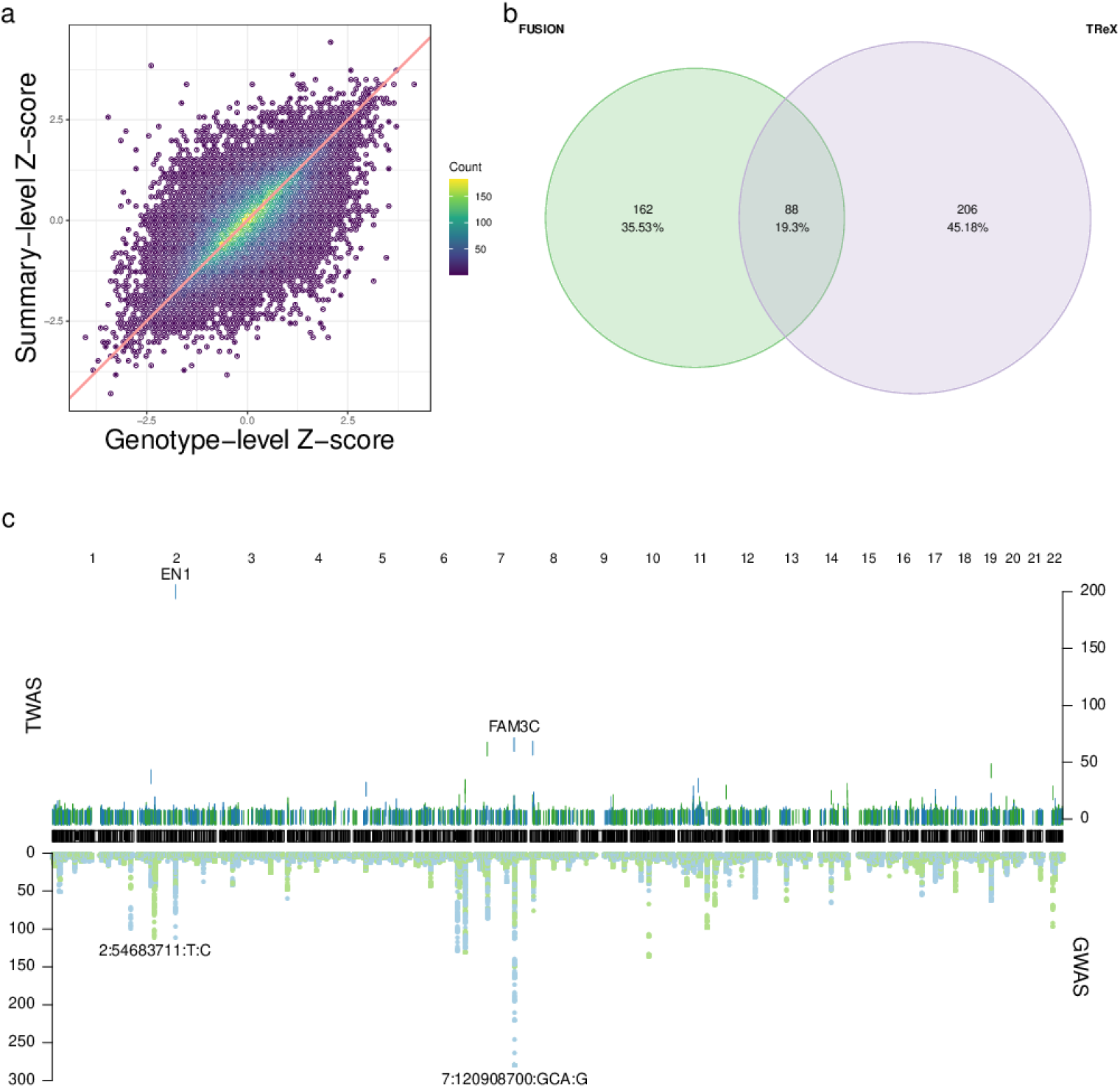
Summary-level TWAS. **a.** Comparison of the genotype- and summary-level TWAS for TReX. In both cases the weights were learned in the brain cortex of GTEx white samples and the phenotypes was the level of amyloid *β* at the baseline visit for white ADNI samples. For the summary-level TWAS, the linkage disequilibrium matrix was obtained from all GTEx white samples. Pearson correlation: 0.66, t = 166.47, df = 36012, p-value *<* 2.2e-16. **b.** Comparison of genes found by FUSION and TReX. The Euler diagram shows the protein-coding genes with an association p-value smaller than 1e-9. The weights were trained in 120 white individuals from GTEx using the expression values from the brain cortex, and the Z-scores are from the height GWAS. **c.** Double Manhattan plot for BMD. The top panel shows the genes whose predicted expression in brain cortex is associated with the bone mineral density; the bottom panel shows the SNPs associated to this trait in the UK Biobank data set.

We thus decided to perform a summary-level TWAS on 11 phenotypes from the UK Biobank, including height, body mass index, bone mineral density, platelet count, and various other blood-related metrics (Suppl. Table 3). We used 1e-9 as the study-wide significance threshold for the p-value; this number accounts for testing about 40,000 genes in 48 tissues for 11 phenotypes.

To determine whether TReX has an advantage over other existing tools, we trained the models for FUSION on the same brain cortex data as TReX, and compared the results. Overall, the Z-scores by FUSION and TReX are quite correlated (Pearson corr. 0.38 for the Z-scores pertaining to the TWAS on height, Suppl. Fig. 4). However, TReX found a considerable number of genes with p-value < 1e-9 that were not significant for FUSION. In the height TWAS, TReX detected even more significant genes than FUSION (Figure 3 **b**).

The complete lists of significant associations between genes and the 11 UK Biobank phenotypes analyzed is available as a supplementary file (Section 4.12). Here we discuss some notable examples. By far, the phenotype where we could detect the greatest number of associations was height, with 3526 unique genes having an association p-value smaller than 1e-9 in at least one tissue, and each association detected on average in about 13 tissues. Moreover, 166 genes showed a p-value smaller than 1e-9 in all 48 tissues. At the opposite end, red blood cell count has the smallest number of associations, with only 125 unique genes showing p-values smaller than the threshold (Table 2).

**Table 2:** Number of significant genes for each phenotype. We performed a summary-level TWAS on 11 phenotypes from the UK Biobank; the table reports, for each phenotype, the number of genes with a p-value smaller than 1e-9 in at least one tissue. The threshold comes from 0.05*/*(40, 000 × 48 × 11), since we tested the association of about 40,000 genes in 48 tissues for 11 phenotypes. The third column reports the number of genes reported by the GWAS catalog for the phenotype.

| Phenotype | Genes in the<br>TReX TWAS | Genes in the<br>GWAS catalog | Genes in<br>common |
| --- | --- | --- | --- |
| height | 3526 | 3469 | 712 |
| platelet count | 2107 | 1282 | 341 |
| eosinophil count | 1454 | 1170 | 239 |
| RBC distr width | 1287 | 1145 | 211 |
| WBC count | 1138 | 1374 | 179 |
| BMI | 1067 | 119 | 25 |
| FVC | 1065 | 732 | 71 |
| FEV1 | 1033 | 114 | 20 |
| BMD | 748 | 72 | 12 |
| mean corpuscular haemoglobin | 502 | 1328 | 56 |
| RBC count | 124 | 1429 | 3 |

UQCC1 is well-known for its association with human height [45]; consistently, we found that the expression of this gene in the frontal cortex of the brain is strongly and negatively associated with height. Similarly, ATOH7 and GH2, both previously reported in the GWAS catalog, were also found to be strongly associated with height in our TWAS. At the same time, we found genes that were not reported in the GWAS catalog, but whose association was supported by functional evidence. One example is FRMD8, which promotes growth factor signaling [46]: its expression in the hypothalamus was positively associated with height. IPP is another promising candidate for further investigations, as its expression is negatively correlated with height; this gene, according to STRING [47], interacts with KLHL28, which in turn is known to be associated with body height.

Bone mineral density (Figure 3 **c** and Suppl. Table 4) was associated with 645 genes, among which EN1 (a homeobox-containing gene) is prominent in that it is the most strongly associated gene to any of the 11 phenotypes. Conditional loss of EN1 is associated with low bone mass in mice [48], and many rare variants within this gene have also been previously associated to bone density [48, 49]. Strikingly, the expression of TP53INP2 is associated to bone density in all 48 GTEx tissues, but it is not reported in the GWAS catalog. This gene promotes autophagy and can act as a transcriptional activator, but we found no mention of its involvement with bone mineral density in the literature.

## 3 Discussion

We developed a new way to predict the genetic component of the expression of a gene from the sequence composition of its proximal regulatory sequence parametrized by its profile of affinities for position weight matrices. The profile of affinities of a regulatory region provides an efficient non-linear encoding of the genotype which allows the regression model to capture a larger fraction of the expression variance compared with regression models directly using the SNP genotypes as their independent variables. Contrary to our initial expectations, the predictive power of TBA profiles is not directly linked to their ability to predict transcription factor binding [18], as scrambled PWMs, which do not represent the binding preferences of actual transcription factors, led to very similar results. Therefore, the tranformation of genotype data into TBA profiles should rather be interpreted as a process of feature expansion, i.e. the creation of new features from non-linear combinations of the original features: Indeed the number of PWMs we use to encode the regulatory regions (640) is much higher than the typical number of variants in the regions we consider (mean 11.79 variants).

A related method to predict the variation of expression across individuals was recently put forward by Shi and collaborators [15], who used the DeepSEA [50] algorithm trained on a lymphoblastoid cell line to predict the effect of variants on the binding of transcription factors to their target sites (identified by ChIP-seq on the same cell line). Such predicted binding changes are then used as independent variables in regularized regression to develop predictive models of gene expression. An important difference with our method is that TF2Exp uses cell-type specific epigenetic information (by using DeepSEA trained on lymphoblastoid cell lines, and ChIP-seq and HiC data derived from the same cells to identify binding sites and assigning them to genes). Therefore our method, in which gene expression in the training data is the only tissue-specific information used, is more readily usable in a variety of tissues for which epigenetic information is not yet available. On the other hand, the relationship between variants and transcription factor binding is much more explicit in TF2Exp, which might thus better capture the causal relationship between alterations of TF binding and gene expression.

It is surprising that our model achieves such good results by just considering ~2,000 bps of proximal regulatory regions, although also variant-based methods such as TIGAR, while considering in principle SNPs in much larger regions, show larger effect sizes for those in the proximity of the TSS, and the same was observed for eQTLs [51]. Limiting the analysis to proximal regions leads to a decrease in both signal and noise, in such a way that their ratio might be actually increased. It should also be kept in mind that the proximal sequences do contain, through linkage disequilibrium, significant information about genetic variants located tens of thousands of base pairs away. Finally, proximal regulatory regions can be assigned to their target genes with much more confidence than distal ones, thus reducing the risk of multiple-hit genes (that is, multiple genes being associated to a phenotype based on the same locus [52]). When we tried to include enhancers in our model we faced two problems, the first being how to associate the enhancers to their target genes in a tissue-specific way, and the second being how to combine the TBAs of proximal and distal regions. As our attempts, so far, did not result in a significant increase in cross-validated *R*^2^ compared with the proximal-only models used here, solving these two problems will be the focus of our next efforts.

Our method is not immune to the general limitations of the TWAS approach, including the inability to distinguish causation from pleiotropy, hence we would still recommend that TWAS be complemented by other methods such as colocalisation and mendelian randomization.

In conclusion, regressing gene expression on regulatory sequence composition parametrized thorugh PWMs leads to improved prediction of the genetic component of gene expression, thus allowing the discovery of new gene/trait associations via TWAS.

## 4 Methods

### 4.1 Genotypes

We obtained individual-level genotypes from three large-scale projects: GTEx v7 [53] (accession number phs000424.v7.p2), GEUVADIS [54], and ADNI (http://adni.loni.usc.edu/). Our method requires phased haplotypes, but not all the VCF files provided by the three above projects were phased. For GTEx, we performed the phasing using Beagle (28Sep18.793) with the HapMap GrCh37 genetic map. For GEUVADIS, the haplotypes were already phased for most of the samples and we discarded those for which they were not. For ADNI, we performed the phasing and imputation using Beagle (18May20.d20) [55, 56], using the 1000 Genomes Project phase 3 as the reference panel and the HapMap GrCh37 genetic map (everything was downloaded from http://faculty.washington.edu/browning/beagle/beagle.html), setting the window to 70.0 cM; the conform-gt program was used beforehand to adjust the positions and allele order of the genetic variants of the raw VCF to match the reference panel, as suggested in Beagle’s manual.

The individual-level, phased genotypic data obtained from the three projects were uniformly processed before performing further analyses. For each data set, the VCF files were filtered using plink2 [57] as follows (the command line options are indicated in brackets). First, the files were separated by population (`--keep`); for this work we focused on white individuals for GTEx, white individuals for ADNI, and Yoruban individuals for GEUVADIS. We removed the genetic variants that failed QC tests (`--var-filter`) or that were duplicated (`--rm-dup`). Individuals with more than 10% of the markers missing were excluded (`--geno 0.1`). Variants with a Hardy-Weinberg p-value (with the “midp” correction) less than 10^−10^ were filtered out (`--hwe 1e-10 midp`). Only biallelic markers (`--max-alleles 2`) with a minor allele frequency of at least 1% (`--maf 0.01`) were retained. Moreover, the variant names were set to a unique, unambiguous identifier (--set-all-var-ids ‘@:#:$r:$a’ --new-id-max-allele-len 100 missing). Finally, the result was exported again in VCF format (`--export id-delim=‘,’ vcf-4.2`)

### 4.2 Total binding affinity

In a 2006 article by Foat *et al.* [17], a statistical-mechanical model was used to derive a formula for the occupancy of a DNA sequence by a transcription factor, motivated by the fact that in many experiments (such as ChIP-chip or differential mRNA expression) the signal is, at least approximately, proportional to the occupancy. Inspired by this work, Molineris *et al.* [19], introduced a similar approach, but with two novelties: (i) PWMs are used to compute the affinity, and (ii) for each possible binding site, the score is given by the maximum between the forward and reverse strand. In particular, the <u>total binding affinity</u> of the transcription factor associated to the PWM *m* for the regulatory region of gene *g* is defined as

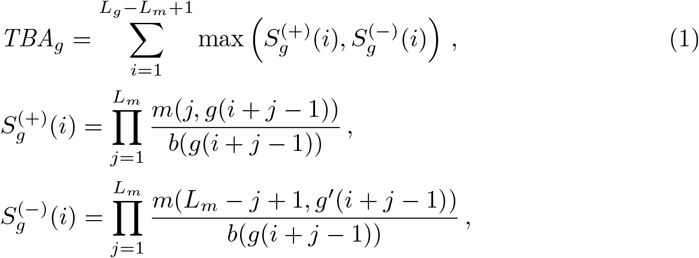

where: 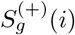 and 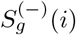 are the scores of motif *m* starting at position *i* of sequence *g* on the forward and reverse strands, respectively; *g*(*i*) is the base at position *i* of the regulatory region *g*; *g*′ (*i*) is the base at the same position but on the opposite strand, i.e. the Crick-Watson complement of *g*(*i*); *m*(*i, g*(*i*)) is the probability of observing base *g*(*i*) at position *i* in the PWM of transcription factor *k*; *b*(*g*(*i*)) is the background probability of observing base *g*(*i*).

The computation of the total binding affinity requires the genotypes in VCF format, the reference DNA sequence of the model organism, a BED file specifying the coordinates of the regulatory regions of each gene, the PWMs of all the transcription factors, and the background nucleotide frequencies. Helper scripts to find the regulatory regions and compute the total binding affinity are provided with this article (Section 4.11). In particular, the script to compute the total binding affinity relies on a specialized program called vcf rider [58]. For this article, the affinity was computed for white individuals in GTEx [10], white individuals in ADNI, and Yoruban individuals in GEUVADIS [54], using the UCSC hg19 sequence as reference.

The regulatory regions associated to a gene were chosen to be sequences spanning (−1500, +500) from the transcription start site of all annotated transcripts of the gene. The annotation of the transcripts was derived as follows. For GTEx and ADNI, we used the reference annotation from the GTEx portal (https://gtexportal.org/home/datasets, downloaded on 07/25/2019), whereas for GEUVADIS we used GENCODE v12 [59]; these annotations contain the coordinates of the transcripts whose expression was measured. If the regulatory regions of two or more transcripts associated to the same gene overlapped, they were merged. After computing the total binding affinity for the set of (non-overlapping) regulatory regions of all the transcripts of a gene, they were summed and log_2_-transformed.

The PWMs were obtained from Hocomoco 10 [20]: we downloaded the mononucleotide models for humans. The background frequencies were calculated on the intergenic regions of the hg19 reference human sequence.

### 4.3 Gene expression

Gene expression data in the form of RPKMs and read counts were obtained from the GTEx [53] and GEUVADIS [54] projects, and were preprocessed as reported in the article describing the analysis of GTEx v7 [53]. Briefly, we kept only the genes with RPKM > 0.1 and read count > 5 in at least 10 samples, we applied a quantile normalization across genes and an inverse-normal transformation across samples, and finally we subtracted the effect of covariates such as the PEER factors [60] (more details below).

Quantile normalization is used to minimize non-biological differences between individuals with respect to the RNA-sequencing [61]. Using the jargon of [61], each individual is treated as a dataset and each gene as an observation. The observed quantiles are projected onto the unit diagonal, and after this operation the distribution of gene expression is the same for all the individuals. An inverse-normal transformation [62] is then applied so that the distribution of the expression of each gene across individuals becomes standard normal. These transformations are applied to expression data for all tissues together, then the expression data are split by tissue.

Before training the models for the TReX imputation expression data were replaced by their residuals after the linear regression *EXPRESSION* ~ *COVARIATES*. The covariates for each tissue were downloaded from the GTEx portal and included: sex, sequencing platform, the first three principal components (PCs) of the genotype, and a variable number of PEER factors depending on the number of samples with available expression for that tissue. The PEER factors were computed on the top 10000 genes and their number was min(15*, Ceiling* (*SampleSize/*5)) if the sample size was < 150, 30 if the sample size was ≥ 150 and < 250, 35 otherwise.

Although the GEUVADIS expression data had already been preprocessed by the authors of the original study, we started from the raw RPKM and read counts and appiled the above procedure also on the GEUVADIS data so as to be able to compare the results across the GTEx and GEUVADIS data sets.

### 4.4 TReX imputation

In the classical approach, where each SNP is encoded as 0, 1, or 2 according to its genotype, the following model is fitted for each gene:

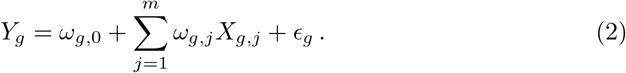

Here, *Y_g_* is the residualized expression of gene *g*, *m* is the total number of markers in a window (tipically 1Mbp) around gene *g*, and *X_g,j_* is the genotype at locus *j*. The intercept *ω*_*g*,0_ captures the baseline expression value for gene *g*, while the error term *ϵ* captures the non-cis-genetic variance of gene expression. Typically, both the expression and the genotypes are centered and scaled, so that the incercept can be ignored.

In our approach, the effect of many SNPs is integrated over the length of the proximal regulatory region of gene *g* through the total binding affinity. The total binding affinity of each PWM captures the effect of all the SNPs simultaneously, although each SNP will have different effects on different PWMs according to which motifs it breaks or creates. Our Ridge regression fits the following model for each gene,

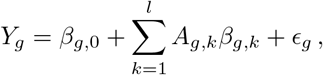

where *l* is the total number of available PWMs (640 for HOCOMOCO v10), and *A_g,j_* is the log_2_-transformed total binding affinity for transcription factor *k*, as outlined in 4.2.

Model performance was evaluated for all tissues by 5-fold nested cross validation, whereby the samples are split in 5 groups and the model is trained 5 times, each time leaving one of the groups out and using the samples in that group to evaluate the performance of the model trained on all the other samples. Furthermore, in order to estimate the best value of the Ridge penalty factor, *λ*, we used a nested 10-fold cross-validation, where each training group is further split in 10 subfolds and each subfold is used in turn to evaluate the performance of a model trained by choosing the parameter from a grid of possible values; this inner cross-validation loop is used to find the optimal value of the parameter *λ* in a way that does not bias the evaluation of the performance that is carried out in the outer loop. For the sake of obtaining the prediction weigths, however, the outer cross-validation loop was omitted and only the inner, 10-fold loop was retained.

The results of the training are *l* weights for each gene, where each weight represent the effect of the affinity of one particular PWM on the expression of that gene. Then, it is possible to compute the total binding affinities of the promoters of each gene in a testing data set for all the individuals, and to use the weights to “predict” the expression values in the testing data set. Software for performing all these tasks is avalable (see Section 4.11).

### 4.5 Phenotypes

For the ADNI data set, we started from the adnimerge table provided by the AD-NIMERGE R package version 1 (https://adni.bitbucket.io/), which contained various phenotypes measured at multiple time points for each individual, during the baseline visit and subsequent follow ups. We kept only individuals annotated as “white” and for which the genotypes were available, resulting in 735 samples, and for them we retained only the values maesured at the baseline visit. Furthermore, we excluded the following phenotypes because we used them as covariates: age (at baseline), sex, years of education, ethnicity, marital status, number of APOE epsilon4 alleles, intracranial volume (at baseline).

The disease status at baseline was originally encoded as a categorical variable with the following values: healthy controls (n = 253), early mild cognitive impairment (n = 211), late mild cognitive impairment (n = 228), significant memory concern (n = 0), Alzheimer’s disease (n = 43). We recoded this variable in two different ways and considered the resulting variables as two distinct phenotypes: in the first, AD samples were encoded as “1” and all the others as “0”, whereas in the second CN samples were encoded as “0” and all the others as “1”. We found that the latter encoding provided the most significant associations, possibly because the two classes are of similar size, whereas in the first encoding, only 5% of the individuals belong to the “1” class. However, when discussing associations with the disease status in the results, we refer to associations with any of these two encodings.

All the other 41 remaining baseline phenotypes were quantitative; examples of these include volumetric measures of various regions of the brain, scores of cognitive tests, and quantifications of amyloid beta and tau proteins. Quantitative phenotypes that ranged outside of the sensitivity intervals (and were therefore denoted like “*< x*” or “*> x*”) were truncated to *x*.

### 4.6 Genotype-level TWAS

Association between the 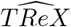 in each tissue and each phenotype was assessed by linear or logistic regression for quantitative and binary phenotypes, respectively. For all the regressions, the formula was 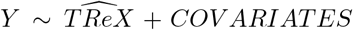, where 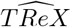 is the imputed expression and the covariates are: age (at baseline), gender, years of education, ethnicity, marital status, number of APOE epsilon4 alleles, intracranial volume (at baseline), and the first ten principal components of the genotype. Aside from the PCs of the genotype, which were computed with plink2 from the genotypes preprocessed as described in Section 4.1, all the other covariates were derived from the adnimerge table from the ADNIMERGE R package. The Z-score and the p-value of the association between expression and phenotype were defined as, respectively, the standardised coefficient and the p-value of the variable 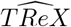 in the regression model. All the software required to perform the genotype-level TWAS is publicly available (Section 4.11).

### 4.7 GWAS

In order to compare the genotype-level with the summary-level TWAS we needed the summary statistics of a GWAS performed on the ADNI data set. We used plink2 to convert the population-specific VCF files preprocessed as described in Section 4.1 into plink’s binary format, and passed to plink2 the phenotypes and covariates pro-cessed as described in Section 4.5. Besides `--glm` and the options for the genotpe, phenotype, and covariates files, the only additional option we passed to plink2 was `--covar-variance-standardize`.

### 4.8 Summary statistics

For the ADNI data set, the output of plink2 consisted of the summary statistics; for the UK Biobank, we downloaded summary statistics for 11 of the 7221 phenotypes which were considered as part of a GWAS analysis across 6 continental ancestry groups by the Neale lab [63]. We processed all the summary statistics first with `fizi munge` and then with fizi impute (https://github.com/bogdanlab/fizi). The former parses the summary statistics and performs some quality control checks before printing the data in a standardized format, while the latter imputes the missing Z-scores.

`fizi` can leverage functional data to improve the imputation, but in this case we did not use this feature, and performed the imputation using only a reference linkage disequilibrium panel. The reference linkage disequilibrium file is simply a plink.bed file with the genotypes of all the individuals in a data set. Here, we derived the reference LD file from the GTEx data set processed as described in Section 4.1, thus including only white individuals. During the imputation procedure, a variant present in the reference file but without a Z-score is assigned a Z-score computed from the scores of its neighbouring variants and the respective linkage disequilibrium in the reference panel.

### 4.9 Summary-level TWAS

In the original approach to a summary-level TWAS [7], the association test relies on the following Wald statistic:

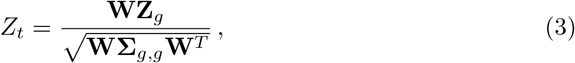

where: *Z_t_* is the TWAS Z-score; **Z**_*g*_ is the vector of GWAS Z-scores of the SNPs within 1Mbp around the gene; **W** is the vector of weights, one for each SNP, obtained by the predictive model; and **Σ**_*g,g*_ is the linkage disequilibrium matrix between the SNPs in the window. In practice, **Σ**_*g,g*_ is unknown because we lack the full genotypes from the GWAS data set, but it can be estimated from a separate reference linkage disequilibrium panel.

This approach requires in particular **W**, the effect of each SNP on gene expression, while we only have the effect of the affinity on gene expression. However, since the total binding affinity is a deterministic function of the genotype (Section 4.2), we can compute the approximate change in total binding affinity caused by each SNP, and from that the change in gene expression caused by each SNP. We computed this quantity empirically as the difference between the mean affinity of all the individuals with genotype 0 and the mean affinity of all the individuals with genotype 1 in a reference data set. Since we had already computed the affinities for all the individuals in GTEx, we used it as the reference for computing what we call the Δ_*TBA*_.

For this article we used GWAS Z-scores either from the GWAS that we performed on ADNI (4.7) or from the UK Biobank; we preprocessed them as described in Section 4.8. Summary results for 11 UK Biobank complex traits were obtained from the Neale Lab web site (http://www.nealelab.is/uk-biobank) and are listed in Suppl. Table 3. The expression weights *β* were trained on white individuals in GTEx, and both the Δ_*TBA*_ and **Σ** were derived from the genotypes of white individuals in GTEx. All the scripts are available on GitHub (Section 4.11).

### 4.10 TIGAR and FUSION

We downloaded the software from GitHub (https://github.com/yanglab-emory/TIGAR and https://github.com/gusevlab/fusion_twas) and followed the authors’ instructions to compute the weights using our own genotype and expression files (the same we used for TReX). For TIGAR, we trained the DPR model and evaluated the 5-fold cross-validation *R*^2^. For FUSION we trained all the available models except BSLMM due to the high computational cost of the latter; in all cases we used the default parameters specified by the authors of the software.

## Supporting information

Supplementary file: ADNI associations

Supplementary file: UKBB associations

## 4.11 URLs

**GitHub repository** https://github.com/fmarotta/TReX

## 4.12 Supplementary files

- TWAS ADNI significant.tsv
- TWAS UKBB significant.tsv

## 4.13 Acknowledgements

The Genotype-Tissue Expression (GTEx) Project was supported by the Common Fund of the Office of the Director of the National Institutes of Health (common-fund. nih.gov/GTEx). Additional funds were provided by the NCI, NHGRI, NHLBI, NIDA, NIMH, and NINDS. Donors were enrolled at Biospecimen Source Sites funded by NCI Leidos Biomedical Research, Inc. subcontracts to the National Disease Research Interchange (10XS170), Roswell Park Cancer Institute (10XS171), and Science Care, Inc. (X10S172). The Laboratory, Data Analysis, and Coordinating Center (LDACC) was funded through a contract (HHSN268201000029C) to the The Broad Institute, Inc. Biorepository operations were funded through a Leidos Biomedical Research, Inc. subcontract to Van Andel Research Institute (10ST1035). Additional data repository and project management were provided by Leidos Biomedical Research, Inc.(HHSN261200800001E). The Brain Bank was supported supplements to University of Miami grant DA006227. Statistical Methods development grants were made to the University of Geneva (MH090941 & MH101814), the University of Chicago (MH090951,MH090937, MH101825, & MH101820), the University of North Carolina - Chapel Hill (MH090936), North Carolina State University (MH101819),Harvard University (MH090948), Stanford University (MH101782),Washington University (MH101810), and to the University of Pennsylvania (MH101822). The datasets used for the analyses described in this manuscript were obtained from dbGaP at http://www.ncbi.nlm.nih.gov/gap through dbGaP accession number phs000424.v7.p2/GRU.

## Notes

### Competing Interest Statement

The authors have declared no competing interest.

### Summary of Updates

Acknowledgement added. Some typos corrected.

