## Supplementary text, figures, and tables for "Prediction of gene expression from regulatory sequence composition enhances transcriptome-wide association studies"

#### 1 Supplementary note

##### 1.1 Derivation of the test statistic for the summary-level TWAS

In the original approach to a summary-level TWAS [**gusev˙2016**], the association test relies on the following Wald statistic:

$$Z_t = \frac{\mathbf{W}\mathbf{Z}_g}{\sqrt{\mathbf{W}\mathbf{\Sigma}_{g,g}\mathbf{W}^T}}\,. \tag{1}$$

As we understood it, the assumption is that, in the absence of any associations, the TWAS Z-score ($Z_t$) of one gene and the vector of GWAS Z-scores ($\mathbf{Z}_g$) of the SNPs around the gene are jointly Gaussian with mean zero. This means that the quantity $Z_t|\mathbf{Z}_g$ is Gaussian with mean $\mu_{t|g} = \mu_t + \mathbf{\Sigma}_{t,g}\mathbf{\Sigma}_{g,g}^{-1}(\mathbf{Z}_g - \mu_g)$, which allows one to estimate the TWAS Z-score as $\bar{Z}_t = \mathbf{\Sigma}_{t,g}\mathbf{\Sigma}_{g,g}^{-1}\mathbf{Z}_g$. Here, $\mathbf{\Sigma}_{g,g}$ is the linkage disequilibrium matrix between the GWAS SNPs and $\mathbf{\Sigma}_{t,g}$ is the covariance between the expression of the gene and the genotypes for the SNPs in the window around the gene. Letting for simplicity $\mathbf{W} = \mathbf{\Sigma}_{t,g}\mathbf{\Sigma}_{g,g}^{-1}$, we estimate $\bar{Z}_t = \mathbf{W}\mathbf{Z}_g$. Now, if the gene expression is imputed with a linear regression model as in Equation (2) of the main text, the vector of coefficient $\omega$ is given by $(\mathbf{X}^T\mathbf{X})^{-1}\mathbf{X}^T\mathbf{Y}$, where $\mathbf{X}$ is a matrix of genotypes and $\mathbf{Y}$ is a vector of gene expressions; moreover, if the genotype and expression vectors are centered, theoretically,

$\mathbf{\Sigma}_{g,g} = (\mathbf{X}^T\mathbf{X})^{-1}$ and $\mathbf{\Sigma}_{t,g} = \mathbf{X}^T\mathbf{Y}$, *i.e.* $\mathbf{W} = \omega^T$. In summary, it is possible to combine the Z-scores from a GWAS with the expression weights to get an estimate for the TWAS Z-score. For the actual association test, we divide by the standard deviation of the estimate and obtain Equation 1.

This approach requires in particular $\mathbf{\Sigma}_{t,g}$, the effect of each SNP on gene expression, while, by using the total binding affinity, we only have the effect of the affinity on gene expression. However, since the total binding affinity can be computed as a known, deterministic function of the genotype (section 4.2), we are able to find the approximate change in total binding affinity caused by each SNP, and from that we can compute the change in gene expression caused by each SNP.

More formally, let $Y$ be the expression of a gene, $\mathbf{X}$ be an $m$-vector of genotypes, and $\mathbf{A}$ be an $l$-vector of $\log_2$-transformed affinities. Rewriting the classical model (Equation 2 of the main text) in vector notation, we have

$$Y = \omega \cdot \mathbf{X}\,, \tag{2}$$

where $\omega$ is an $m$-vector of weights trained in an eQTL data set, whereas in our model $Y = \beta \cdot \mathbf{A}$, where $\beta$ is an $l$-vector. The total binding affinity is a function of the genotype, so if we imagine that the affinity be a continuous, approximately linear function of a continuous genotype (as the blue line in Figure 5, left panel), we can write $\mathbf{A} = \mathbf{g}(\mathbf{X})$ and, substituting in Equation (2),

$$Y = \beta \cdot \mathbf{g}(\mathbf{X})\,. \tag{3}$$

To find $\omega$ from Equation (2), we would compute the (multivariable) derivative with respect to $\mathbf{X}$; similarly, to find the change in expression attributable to a change of one unit in the genotype, we can compute the derivative of Equation 3 with respect to $\mathbf{X}$. Using the chain rule from multivariable calculus, the weight for SNP $i$ on gene expression is given by

$$W_i = \frac{dY}{d\mathbf{A}} \cdot \frac{d\mathbf{A}}{dX_i}\,. \tag{4}$$

In practice, $\frac{dY}{d\mathbf{A}}$ is the vector $\beta$ and $\frac{d\mathbf{A}}{dX_i}$ is the vector whose generic element is $\frac{dA_j}{dX_i}$, *i. e.* the change in the $\log_2$ of total binding affinity of transcription factor $j$ attributable to a change of 1 in the genotype of SNP $i$. Computing the derivative of the $\log_2$ of the formula for the TBA is an impossible task because, first of all, the genotype is discrete, and secondly, the TBA depends also on the reference sequence, not just on the 0/1/2-encoded genotype. Therefore, we computed $\frac{d\mathbf{A_j}}{dX_i}$ empirically as the difference between the mean $A_j$ of all the individuals with genotype 0 and the mean $A_j$ of all the individuals with genotype 1 in a reference data set. Equation 4 allows us to find $\mathbf{W}$, which we then plug into Equation 1 to obtain the TWAS Z-score. This chain-rule approach can be used also with other functions, if more efficient predictors than the total binding affinities should be discovered in the future.

#### 1.2 A possible example of reverse causality

In the main text we discuss the anecdotal case of MAF1, a gene whose predicted expression in several tissue, but not the hippocampus, is strongly associated to the volume of the hippocampus. One possibility to explain this and similar cases is with reverse causality.

We illustrate this argument with a small thought experiment. Suppose there is a genetic variant which increases the expression of MAF1 in all the tissues where the gene is expressed; suppose also that in the hippocampus (and only in the hippocampus) a high expression of MAF1 leads to a small volume of this brain region, and, in turn, a small hippocampus decreases the expression of MAF1. Thus, in the training data set, individuals with such genetic variant will have a high expression of MAF1 in, say, the brain cortex, a small hippocampus, and a low expression of MAF1 in the hippocampus (because of the feedback). On the other hand, individuals without the genetic variant will have a low expression of MAF1 in the brain cortex, a large hippocampus, and a small expression of MAF1 in the hippocampus (because they do not have the genetic variant). Therefore, individuals with either a large or a small hippocampus will have small expression values for MAF1 in the hippocampus, but this is not true for the brain cortex, where the relationship between the expression of MAF1 and the volume of the hippocampus can be seen because it is not masked by the effect that the phenotype has on the expression in the hippocampus.

#### 1.3 The effect of a variant depends on the others

The total binding affinity can, at least in part, capture long-range relationships between genetic variants. This is because of the $\log_2$ transformation that we apply to the total binding affinities. Suppose that SNP "A" increases the non-transformed total binding affinity by 10 units in an individual whose promoter differs from the reference sequence only at that variant. In another individual, whose promoter differs from the reference seqence at the SNP "A" *and* at SNP "B", the SNP "A" increases the non-transformed affinity by 10 and SNP "B" increases the affinity by, say, 20. After the $\log_2$ transformation, SNP "A" will give different contribution to the total binding affinities in the two individuals.

This introduces some non-additivity which violates the assumptions of the summary-level TWAS; however, the way we compute the $\Delta_{TBA}$ relies on the locally linear relationsip between genotype and total binding affinity, as if we approximated the total binding affinity with its first order Taylor expansion.

#### 2 Supplementary figures

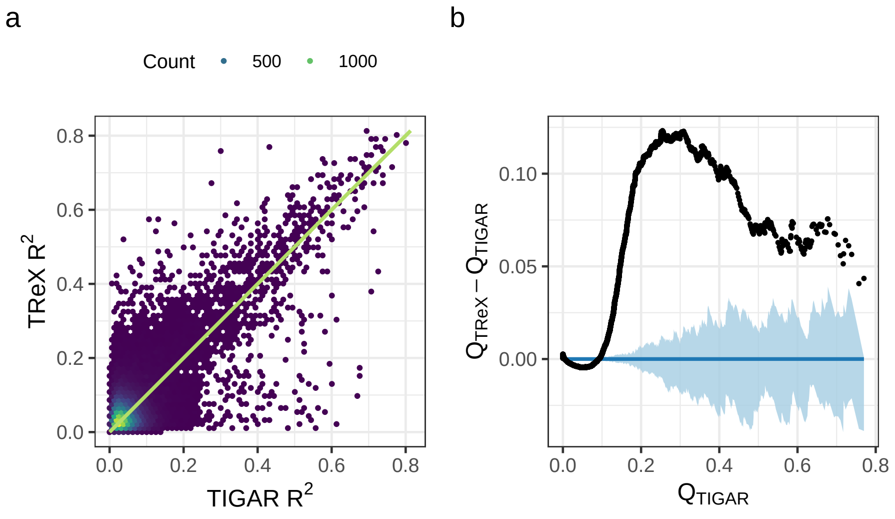


Suppl. Fig. 1: Predictive performance of various models. **a.** Scatter plot of the 5-fold cross-validation $R^2$ for TIGAR and TReX on brain cortex; the plot shows 39770 genes. In principle each point represents a gene, but in order to reduce overplotting we associated potentially many genes that fell approximately in the same region to a single point; the color of a point denotes the count of genes associated to it. The green line is the identity line. **b.** Detrended QQ-plot of the 5-fold cross-validation $R^2$ for TIGAR and TReX on brain cortex. On the x axis there are the quantiles from TIGAR's $R^2$, while on the y axis there are the differences between corresponding quantiles from TReX's and TIGAR's $R^2$. The confidence interval (light blue) was generated from 100 bootstrap replicates from the distribution of TIGAR's $R^2$.

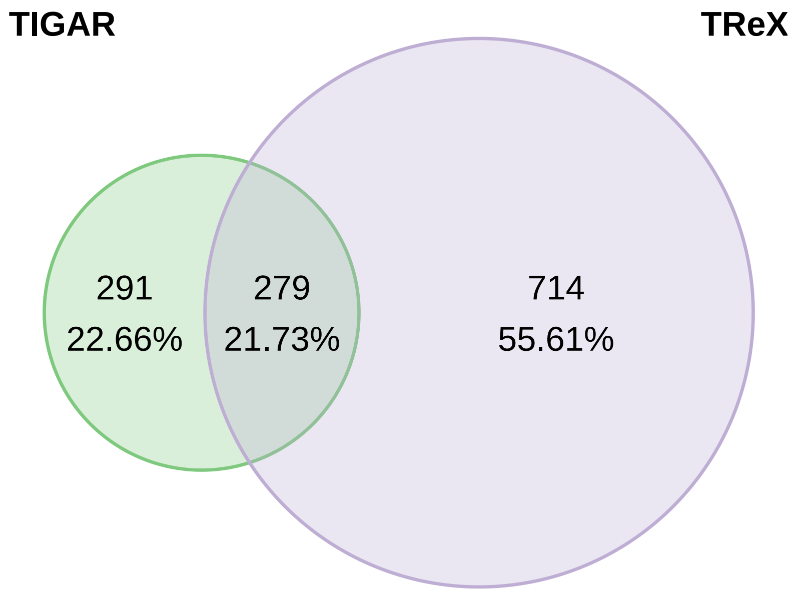


Suppl. Fig. 2: Comparison of the performances of various tools in an external dataset. The figure shows the number of genes with $R^2 > 0.1$ for TIGAR and TReX (among a total of 21567 genes common to both). The models were trained on EBV-transformed lymphocytes from GTEx and applied to Yoruban individuals from GEUVADIS.

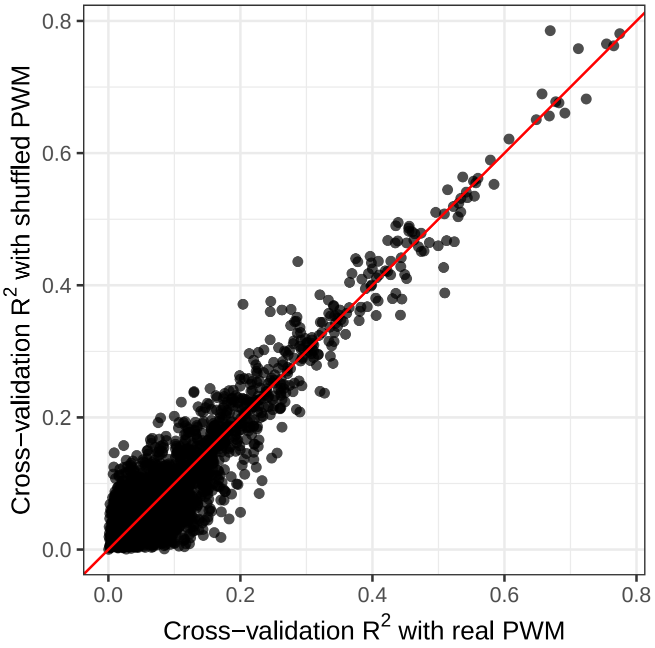


Suppl. Fig. 3: Performance with real and shuffled PWMs. The Cross-validation $R^2$ in brain cortex from GTEx are virtually the same. The red line is the identity line.

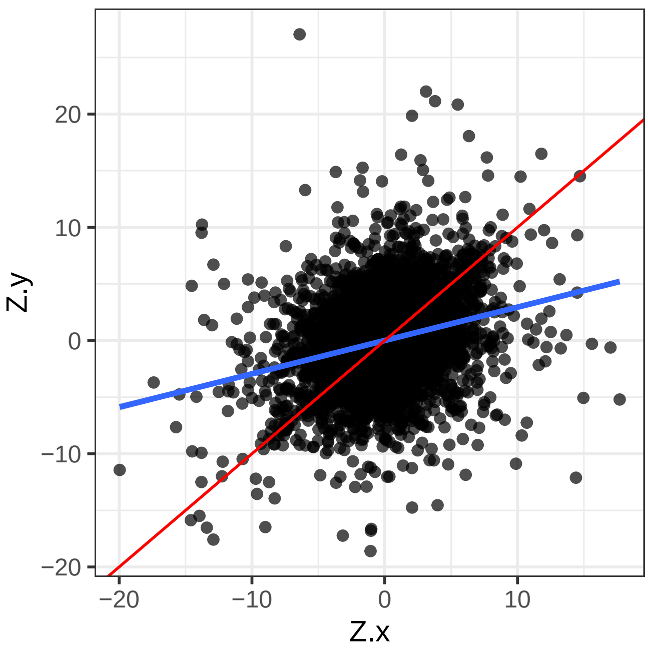


Suppl. Fig. 4: Scatter plot of FUSION and TReX Z-scores. The model was trained on brain cortex and used to perform a TWAS on height. The red line is the identity line, while the blue line is the linear regression line.

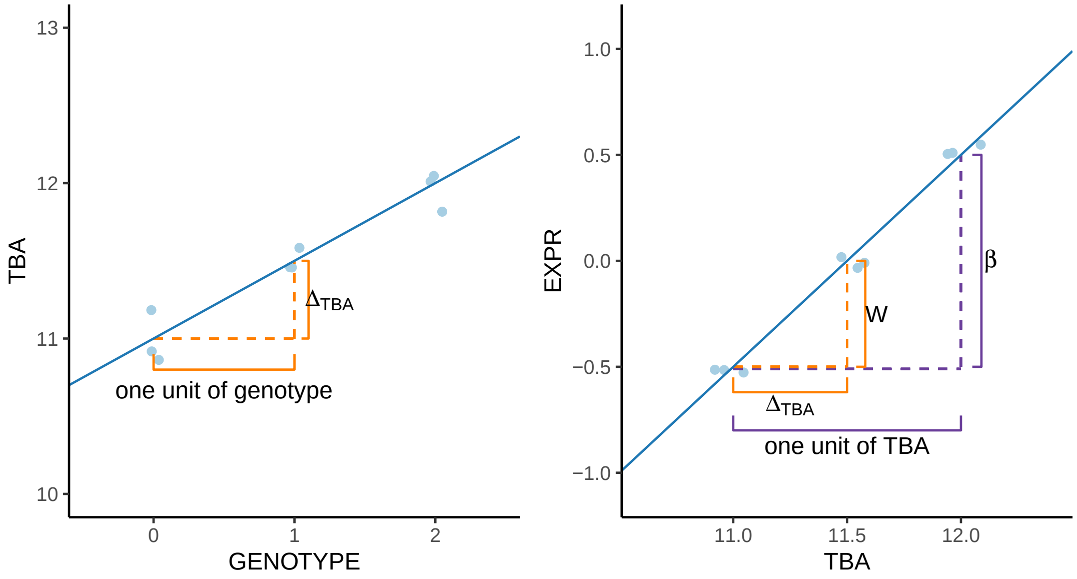


Suppl. Fig. 5: Schematic of $\Delta_{TBA}$. A change of "one unit of genotype" results in a change of $\Delta_{TBA}$ units of the total binding affinity (left panel). A change of one unit of total binding affinity results in a change of $\beta$ units of predicted expression (right panel). And a change of $\Delta_{TBA}$ units of total binding affinity results in a change of $W$ units of predicted expression.

### 3 Supplementary tables

Suppl. Table 1: Cross-validation $R^2$ for all tissues. The table reports, for each tissue in GTEx, the average $R^2$ across all genes in that tissue.

| Tissue | Average $R^2$ |
|---|---|
| Brain_Substantia_nigra | 0.088 |
| Minor_Salivary_Gland | 0.086 |
| Brain_Spinal_cord_cervical_c | 0.084 |
| Uterus | 0.079 |
| Brain_Amygdala | 0.078 |
| Cells_EBV_transformed_lymphocytes | 0.071 |
| Brain_Putamen_basal_ganglia | 0.070 |
| Vagina | 0.070 |
| Brain_Cerebellar_Hemisphere | 0.069 |
| Ovary | 0.069 |
| Brain_Frontal_Cortex_BA | 0.068 |
| Brain_Anterior_cingulate_cortex_BA | 0.068 |
| Brain_Hypothalamus | 0.067 |
| Spleen | 0.067 |
| Brain_Hippocampus | 0.067 |
| Small_Intestine_Terminal_Ileum | 0.067 |
| Brain_Cerebellum | 0.065 |
| Brain_Cortex | 0.065 |
| Brain_Nucleus_accumbens_basal_ganglia | 0.063 |
| Prostate | 0.061 |
| Brain_Caudate_basal_ganglia | 0.060 |
| Testis | 0.059 |
| Artery_Coronary | 0.057 |
| Pituitary | 0.054 |
| Liver | 0.052 |
| Pancreas | 0.051 |
| Adrenal_Gland | 0.050 |
| Nerve_Tibial | 0.049 |
| Esophagus_Gastroesophageal_Junction | 0.049 |
| Colon_Sigmoid | 0.049 |
| Cells_Transformed_fibroblasts | 0.048 |
| Thyroid | 0.047 |
| Artery_Aorta | 0.046 |
| Esophagus_Muscularis | 0.045 |
| Colon_Transverse | 0.045 |
| Breast_Mammary_Tissue | 0.044 |
| Heart_Atrial_Appendage | 0.043 |
| Stomach | 0.043 |
| Esophagus_Mucosa | 0.043 |
| Skin_Not_Sun_Exposed_Suprapubic | 0.043 |
| Adipose_Subcutaneous | 0.042 |
| Artery_Tibial | 0.042 |
| Skin_Sun_Exposed_Lower_leg | 0.042 |
| Adipose_Visceral_Omentum | 0.040 |
| Heart_Left_Ventricle | 0.040 |
| Lung | 0.039 |
| Whole_Blood | 0.034 |
| Muscle_Skeletal | 0.033 |

Suppl. Table 2: Genotype-level TWAS in the ADNI data set. The table displays all the genes with a p-value $< 0.05$ for the association with the phenotypes from the ADNI data set, after applying a Bonferroni correction within each tissue/phenotype pair.

| Gene name | Phenotype | Tissue |
|---|---|---|
| AC006126.4 | EcogPtLang.bl | Heart_Atrial_Appendage |
| AC006130.3 | CDRSB.bl | Brain_Frontal_Cortex_BA |
| AC011247.3 | RAVLT.forgetting.bl | Prostate |
| AC016697.6 | TAU.bl | Brain_Amygdala |
| AC016697.6 | PTAU.bl | Thyroid |
| AC016697.6 | TAU.bl | Thyroid |
| AC092574.1 | PTAU.bl | Adrenal_Gland |
| AC092574.1 | TAU.bl | Adrenal_Gland |
| AC114788.2 | MOCA.bl | Muscle_Skeletal |
| AGTPBP1 | WholeBrain.bl | Ovary |
| ANKRD50 | LDELTOTAL.bl | Brain_Spinal_cord_cervical_c |
| ARRDC5 | Ventricles.bl | Testis |
| CALM1P2 | EcogSPMem.bl | Testis |
| CAPN15 | DX.bl_CN_vs_ALL | Colon_Transverse |
| CTB-51J22.1 | RAVLT.learning.bl | Brain_Hippocampus |
| CTB-51J22.1 | RAVLT.learning.bl | Brain_Substantia_nigra |
| CTB-51J22.1 | RAVLT.learning.bl | Nerve_Tibial |
| CTB-51J22.1 | RAVLT.learning.bl | Skin_Not_Sun_Exposed_Suprapubic |
| CTB-51J22.1 | RAVLT.learning.bl | Vagina |
| CTC-459F4.8 | EcogPtMem.bl | Brain_Spinal_cord_cervical_c |
| CTC-492K19.4 | EcogPtVisspat.bl | Ovary |
| CTD-2524L6.3 | MOCA.bl | Adipose_Visceral_Omentum |
| CTD-2524L6.3 | MOCA.bl | Artery_Tibial |
| CTD-2524L6.3 | MOCA.bl | Muscle_Skeletal |
| DIO1 | EcogSPDivatt.bl | Adrenal_Gland |
| DNM3OS | Fusiform.bl | Brain_Hippocampus |
| DNM3OS | Fusiform.bl | Brain_Hypothalamus |
| DNM3OS | Fusiform.bl | Brain_Spinal_cord_cervical_c |
| DNM3OS | Fusiform.bl | Breast_Mammary_Tissue |
| DNM3OS | Fusiform.bl | Minor_Salivary_Gland |
| DNM3OS | Fusiform.bl | Spleen |
| ECHS1 | FAQ.bl | Artery_Coronary |
| ECHS1 | FAQ.bl | Esophagus_Muscularis |
| EHMT2-AS1 | FAQ.bl | Vagina |
| EOGT | RAVLT.learning.bl | Brain_Cortex |
| EOGT | RAVLT.learning.bl | Brain_Frontal_Cortex_BA |
| EOGT | RAVLT.learning.bl | Brain_Hippocampus |
| EOGT | RAVLT.learning.bl | Brain_Hypothalamus |
| EOGT | RAVLT.learning.bl | Skin_Not_Sun_Exposed_Suprapubic |
| EOGT | RAVLT.learning.bl | Testis |
| FAM154A | EcogPtTotal.bl | Ovary |
| FAM154A | EcogPtVisspat.bl | Ovary |
| FIP1L1 | MOCA.bl | Heart_Atrial_Appendage |
| GS1-259H13.7 | FAQ.bl | Brain_Frontal_Cortex_BA |
| GS1-259H13.7 | FAQ.bl | Brain_Hypothalamus |

*Continued on next page*

Suppl. Table 2 – *Continued from previous page*

| Gene name | Phenotype | Tissue |
|---|---|---|
| HBA1 | WholeBrain.bl | Esophagus_Gastroesophageal_Junction |
| HLA-L | RAVLT.perc.forgetting.bl | Artery_Tibial |
| IGKV2-23 | FAQ.bl | Skin_Sun_Exposed_Lower_leg |
| ITGA3 | MMSE.bl | Esophagus_Muscularis |
| ITGA3 | MMSE.bl | Skin_Not_Sun_Exposed_Suprapubic |
| KB-1683C8.1 | EcogPtOrgan.bl | Brain_Cerebellum |
| KLK8 | MMSE.bl | Nerve_Tibial |
| LDHAP5 | EcogPtVisspat.bl | Brain_Cerebellar_Hemisphere |
| MAF1 | Hippocampus.bl | Brain_Cortex |
| MAF1 | Hippocampus.bl | Brain_Spinal_cord_cervical_c |
| MAF1 | Hippocampus.bl | Esophagus_Gastroesophageal_Junction |
| MIR199A2 | Fusiform.bl | Artery_Tibial |
| MIR199A2 | Fusiform.bl | Brain_Amygdala |
| MIR3605 | FAQ.bl | Testis |
| MTMR12 | FAQ.bl | Uterus |
| MTRNR2L8 | CDRSB.bl | Pituitary |
| MTRNR2L8 | CDRSB.bl | Uterus |
| MXD1 | EcogPtVisspat.bl | Vagina |
| MYH16 | FAQ.bl | Esophagus_Mucosa |
| NKX3-1 | RAVLT.immediate.bl | Muscle_Skeletal |
| OR10H4 | CDRSB.bl | Esophagus_Muscularis |
| OR14A16 | MOCA.bl | Adipose_Subcutaneous |
| OR14A16 | MOCA.bl | Minor_Salivary_Gland |
| PGAM1P10 | FAQ.bl | Adipose_Subcutaneous |
| PGAM1P10 | FAQ.bl | Brain_Hippocampus |
| PGAM1P10 | FAQ.bl | Cells_EBV_transformed_lymphocytes |
| PGAM1P10 | FAQ.bl | Small_Intestine_Terminal_Ileum |
| PLXDC1 | FDG.bl | Brain_Hypothalamus |
| PMP22 | TAU.bl | Colon_Transverse |
| PTK2B | RAVLT.immediate.bl | Whole_Blood |
| PTPRM | MidTemp.bl | Vagina |
| RBM43P1 | DX.bl_AD_vs_ALL | Ovary |
| RGS2 | TRABSCOR.bl | Skin_Sun_Exposed_Lower_leg |
| RN7SL250P | ADAS11.bl | Artery_Aorta |
| RN7SL250P | ADAS13.bl | Artery_Aorta |
| RN7SL28P | TRABSCOR.bl | Brain_Amygdala |
| RN7SL28P | TRABSCOR.bl | Brain_Anterior_cingulate_cortex_BA |
| RN7SL28P | TRABSCOR.bl | Pituitary |
| RNU2-39P | Ventricles.bl | Brain_Cerebellar_Hemisphere |
| RNU2-39P | Ventricles.bl | Nerve_Tibial |
| RNU2-39P | Ventricles.bl | Thyroid |
| RP11-1084J3.1 | EcogPtVisspat.bl | Pancreas |
| RP11-1084J3.1 | EcogPtVisspat.bl | Testis |
| RP11-180I4.2 | TRABSCOR.bl | Skin_Sun_Exposed_Lower_leg |
| RP11-184A2.2 | ADAS11.bl | Brain_Spinal_cord_cervical_c |
| RP11-184A2.2 | ADAS13.bl | Brain_Spinal_cord_cervical_c |
| RP11-184A2.2 | MOCA.bl | Brain_Spinal_cord_cervical_c |
| RP11-203E8.1 | EcogPtOrgan.bl | Adipose_Subcutaneous |
| RP11-203E8.1 | EcogPtOrgan.bl | Artery_Tibial |
| RP11-203E8.1 | EcogPtOrgan.bl | Brain_Cerebellar_Hemisphere |

*Continued on next page*

Suppl. Table 2 – *Continued from previous page*

| Gene name | Phenotype | Tissue |
|---|---|---|
| RP11-203E8.1 | EcogPtOrgan.bl | Brain_Cerebellum |
| RP11-203E8.1 | EcogPtOrgan.bl | Brain_Cortex |
| RP11-203E8.1 | EcogPtOrgan.bl | Brain_Frontal_Cortex_BA |
| RP11-203E8.1 | EcogPtOrgan.bl | Brain_Hypothalamus |
| RP11-203E8.1 | EcogPtOrgan.bl | Brain_Nucleus_accumbens_basal_ganglia |
| RP11-203E8.1 | EcogPtOrgan.bl | Brain_Putamen_basal_ganglia |
| RP11-203E8.1 | EcogPtOrgan.bl | Cells_Transformed_fibroblasts |
| RP11-203E8.1 | EcogPtOrgan.bl | Colon_Transverse |
| RP11-203E8.1 | EcogPtOrgan.bl | Esophagus_Mucosa |
| RP11-203E8.1 | EcogPtOrgan.bl | Liver |
| RP11-203E8.1 | EcogPtOrgan.bl | Lung |
| RP11-203E8.1 | EcogPtOrgan.bl | Minor_Salivary_Gland |
| RP11-203E8.1 | EcogPtOrgan.bl | Ovary |
| RP11-203E8.1 | EcogPtOrgan.bl | Pancreas |
| RP11-203E8.1 | EcogPtOrgan.bl | Pituitary |
| RP11-203E8.1 | EcogPtOrgan.bl | Prostate |
| RP11-203E8.1 | EcogPtOrgan.bl | Skin_Sun_Exposed_Lower_leg |
| RP11-203E8.1 | EcogPtOrgan.bl | Spleen |
| RP11-203E8.1 | EcogPtOrgan.bl | Testis |
| RP11-203E8.1 | EcogPtOrgan.bl | Uterus |
| RP11-203E8.1 | EcogPtOrgan.bl | Whole_Blood |
| RP11-361F15.2 | EcogPtOrgan.bl | Heart_Left_Ventricle |
| RP11-361F15.2 | EcogPtOrgan.bl | Uterus |
| RP11-369C8.1 | RAVLT.perc.forgetting.bl | Vagina |
| RP11-376N17.4 | MOCA.bl | Liver |
| RP11-388N2.1 | EcogPtTotal.bl | Brain_Anterior_cingulate_cortex_BA |
| RP11-417L19.2 | AV45.bl | Adrenal_Gland |
| RP11-424M22.3 | MOCA.bl | Artery_Coronary |
| RP11-424M22.3 | MOCA.bl | Brain_Cerebellum |
| RP11-424M22.3 | MOCA.bl | Brain_Spinal_cord_cervical_c |
| RP11-424M22.3 | MOCA.bl | Nerve_Tibial |
| RP11-424M22.3 | MOCA.bl | Stomach |
| RP11-472N13.2 | Ventricles.bl | Brain_Hypothalamus |
| RP11-480I12.10 | DX.bl_AD_vs_ALL | Adrenal_Gland |
| RP11-509J21.3 | TRABSCOR.bl | Brain_Spinal_cord_cervical_c |
| RP11-51B23.3 | EcogPtVisspat.bl | Lung |
| RP11-548K12.10 | RAVLT.perc.forgetting.bl | Liver |
| RP11-58O3.2 | TRABSCOR.bl | Brain_Anterior_cingulate_cortex_BA |
| RP11-657O9.1 | EcogSPVisspat.bl | Adipose_Visceral_Omentum |
| RP11-657O9.1 | EcogSPVisspat.bl | Brain_Caudate_basal_ganglia |
| RP11-657O9.1 | EcogSPVisspat.bl | Brain_Cerebellum |
| RP11-657O9.1 | EcogSPVisspat.bl | Brain_Putamen_basal_ganglia |
| RP11-657O9.1 | EcogSPVisspat.bl | Cells_EBV_transformed_lymphocytes |
| RP11-657O9.1 | EcogSPVisspat.bl | Esophagus_Muscularis |
| RP11-657O9.1 | EcogSPVisspat.bl | Heart_Atrial_Appendage |
| RP11-657O9.1 | EcogSPVisspat.bl | Nerve_Tibial |
| RP11-657O9.1 | EcogSPVisspat.bl | Pituitary |
| RP11-657O9.1 | EcogSPVisspat.bl | Prostate |
| RP11-657O9.1 | EcogSPVisspat.bl | Small_Intestine_Terminal_Ileum |
| RP11-657O9.1 | EcogSPVisspat.bl | Spleen |

*Continued on next page*

Suppl. Table 2 – *Continued from previous page*

| Gene name | Phenotype | Tissue |
|---|---|---|
| RP11-657O9.1 | EcogSPVisspat.bl | Stomach |
| RP11-657O9.1 | EcogSPVisspat.bl | Uterus |
| RP11-657O9.1 | EcogSPVisspat.bl | Vagina |
| RP11-660M5.1 | EcogPtOrgan.bl | Muscle_Skeletal |
| RP11-666F17.2 | FAQ.bl | Brain_Cerebellar_Hemisphere |
| RP11-666F17.2 | FAQ.bl | Cells_Transformed_fibroblasts |
| RP11-685B14.3 | ADAS11.bl | Brain_Hypothalamus |
| RP11-806J6.1 | WholeBrain.bl | Adipose_Visceral_Omentum |
| RP11-806J6.1 | WholeBrain.bl | Adrenal_Gland |
| RP11-806J6.1 | WholeBrain.bl | Breast_Mammary_Tissue |
| RP11-806J6.1 | WholeBrain.bl | Liver |
| RP11-806J6.1 | WholeBrain.bl | Spleen |
| RP11-806J6.1 | WholeBrain.bl | Whole_Blood |
| RP11-95G17.2 | FAQ.bl | Brain_Hypothalamus |
| RP11-95G17.2 | FAQ.bl | Ovary |
| RP3-333B15.4 | ADAS13.bl | Esophagus_Mucosa |
| RP5-823G15.5 | TRABSCOR.bl | Brain_Cortex |
| RP5-823G15.5 | TRABSCOR.bl | Brain_Nucleus_accumbens_basal_ganglia |
| RPL21P135 | ADAS11.bl | Thyroid |
| RPL7AP64 | EcogPtTotal.bl | Brain_Caudate_basal_ganglia |
| TMEM71 | Hippocampus.bl | Esophagus_Muscularis |
| TMIGD1 | FAQ.bl | Stomach |
| TMIGD1 | mPACCdigit.bl | Stomach |
| TRPC7 | ABETA.bl | Brain_Caudate_basal_ganglia |
| TUSC2 | EcogPtOrgan.bl | Brain_Anterior_cingulate_cortex_BA |
| UHRF1 | Ventricles.bl | Artery_Coronary |
| UHRF1 | Ventricles.bl | Heart_Atrial_Appendage |
| WFDC12 | Ventricles.bl | Nerve_Tibial |
| WIF1 | EcogPtTotal.bl | Esophagus_Mucosa |
| WIF1 | EcogSPVisspat.bl | Small_Intestine_Terminal_Ileum |
| WIF1 | EcogPtLang.bl | Testis |
| YIPF1 | EcogSPDivatt.bl | Brain_Caudate_basal_ganglia |
| YIPF1 | EcogSPDivatt.bl | Nerve_Tibial |
| YIPF1 | EcogSPDivatt.bl | Prostate |

Suppl. Table 3: Phenotypes from the UK Biobank. We downloaded the summary statistics for these complex traits and used them to perform summary-level transcriptome-wide association studies.

| Phenotype | ID |
|---|---|
| BMD | 3148_raw |
| BMI | 21001_raw |
| FEV1 | 20150_raw |
| FVC | 3062_raw |
| RBC_count | 30170 |
| RBC_distr_width | 30070_raw |
| WBC_count | 30000_raw |
| eosinophill_count | 30150 |
| height | 50_raw |
| platelet_count | 30080_raw |
| mean_corpuscular_haemoglobin | 30060_raw |

Suppl. Table 4: Top genes associated to BMD.

| Name | Z-score | Tissue |
|---|---|---|
| EN1 | 162.804 | Brain_Cortex |
| CKB | 39.169 | Esophagus_Gastroesophageal_Junction |
| C11orf48 | -31.034 | Brain_Anterior_cingulate_cortex_BA |
| C11orf49 | 28.570 | Brain_Hippocampus |
| WNT16 | 27.615 | Adipose_Subcutaneous |
| RSPO3 | -27.246 | Heart_Atrial_Appendage |
| C11orf48 | -26.153 | Brain_Hypothalamus |
| SGK223 | 23.686 | Brain_Anterior_cingulate_cortex_BA |
| C11orf49 | 22.797 | Brain_Hypothalamus |
| PPP6R3 | 22.546 | Minor_Salivary_Gland |
